# Primed acquisition and microhomology-mediated end-joining cooperate to confer specific CRISPR immunity against invasive genetic elements

**DOI:** 10.1101/831206

**Authors:** Xiaojie Wang, Bo Wu, Zhufeng Zhang, Tao Liu, Yingjun Li, Guoquan Hu, Mingxiong He, Nan Peng

**Author notes:** These authors contributed equally to this work (Xiaojie Wang, Bo Wu).

## Abstract

CRISPR-Cas systems provide archaea and bacteria with adaptive immunity against invasive genetic elements through acquisition of invader-derived spacers. *De novo* spacer acquisition generally adapts spacers from both invaders and hosts with a bias towards invaders, while primed acquisition shows higher specificity to adapt spacers from invaders. Here, in *Zymomonas mobilis* subtype I-F system, primed acquisition showed much higher efficiency than *de novo* acquisition in this system. However, both routes recognised a large proportion of protospacers with the less conserved 5’-end CC PAM. Moreover, primed acquisition showed a preference towards protospacers located at the opposite strand and 3’ direction of the priming protospacer sites, differing from the canonical subtype I-F system. Further, self-spacers were adapted at a higher frequency during *de novo* spacer acquisition, probably leading to self-interference. Importantly, this species employed microhomology-mediated end-joining (MMEJ) for repair of host DNA breaks guided by self-targeted spacers, and overexpression of a host NAD^+^-dependent Ligase-A significantly increased the repair efficiency. In summary, our findings demonstrate that *Z. mobilis* uses primed acquisition for higher specific uptake of invader DNA and employs MMEJ to repair host DNA breaks guided by self-targeted spacers, showing specific immunity against invasive genetic elements.

---

CRISPR-Cas systems, consisting of CRISPR arrays containing identical repeats separated by unique spacers and associated *cas* genes, defend Bacteria and Archaea against invasive plasmids and viruses ^1–3^. CRISPR–Cas adaptive immunity occurs in three stages: acquisition of *de novo* spacers, crRNA biogenesis, and nucleic acid targeting and cleavage ^2,4^. CRISPR-Cas systems acquire DNA fragments of invading nucleic acids (prespacers) and integrate them into CRISPR arrays as spacers, thus forming hereditable immunological memory ^5^. DNA motifs in invasive genetic elements, or in CRISPR loci, play crucial roles in the spacer-acquisition process. For example, the protospacer-adjacent motif (PAM) associated with protospacers ^6–10^ and the leader-repeat junction ^10,11^ have been found to direct spacer acquisition in different CRISPR-Cas systems. Recently, it was found that CRISPR spacer acquisition requires genome-stability proteins ^12,13^ as well as *trans*- and *cis*-acting factors that maintain integration specificity ^14,15^. In a *de novo* spacer acquisition assay, *E. coli* subtype I-E Cas1 and Cas2 have been found to be required ^10,16–18^. Furthermore, the target DNA interference complex is required in the primed spacer acquisition process, which is triggered by a pre-existing spacer matching the target DNA ^16,17,19–21^.

Discrimination of self and non-self DNA is crucial for immunity. CRISPR-Cas systems have developed diverse mechanisms to avoid autoimmunity, including a PAM requirement for target interference ^22,23^. However, because *de novo* spacer acquisition adapts new spacers from host genomic DNA ^9,24^, avoiding host DNA sampling at the CRISPR spacer acquisition stage is more important. *E. coli* has developed a strategy to limit prespacer supply from the host through *Chi* site-assisted reduction of DNA resection in its genome ^25^. Further, primed adaptation allows cells to effectively counter viruses and plasmids that escape CRISPR interference ^16,26–28^, leading to highly efficient acquisition of new *cis*-located spacers of the protospacer of the invading genetic elements with mutations ^7,21,29,30^. Primed adaptation requires functional Cas1, Cas2, and Cas3 proteins, suggesting a functional link between CRISPR interference and primed adaptation ^20,31,32^.

However, adaptation of host self DNA cannot be avoided. In *de novo* spacer acquisition process, 32%, 7%, 16%, and 22.8% or 1.8% (for induction or non-induction of Cas1, Cas2 expression, respectively) of new spacers have been derived from host genomic DNA in *S. thermophilus* subtype II-A ^24^, *S. islandicus* subtype I-A^9^, *Pectobacterium atrosepticum* subtype I-F ^33^, and *E. coli* subtype I-E ^25^ systems. Even in primed spacer acquisition processes, 0.01–0.03% of new spacers have been derived from host genomic DNA in *E. coli* subtype I-E ^30^ and *P. atrosepticum* subtype I-F ^33^ systems, suggesting that host cells might employ different mechanisms to avoid self-spacer guided interference. Here, we studied CRISPR spacer acquisition and interference of the *Zymomonas mobilis* subtype I-F system, to demonstrate that *Z. mobilis* employs primed spacer acquisition for specific uptake of invader DNA and utilizes the MMEJ system to maintain genome stability after self DNA interference.

## Results

### Identification of leaders and PAM variants for primed spacer acquisition

*Z. mobilis* ZM4 encodes a subtype I-F CRISPR-Cas system, including three CRISPR arrays and an adaptation and a DNA interference modules (Fig. 1a). Here, we investigated the effect of leader sequences and PAM variants on primed acquisition in *Z. mobilis*. Plasmids constructed with “NN-S1” cassettes were transformed into *Z. mobilis* ZM4mrr (Fig. 1b). The NC or CN PAM (N = A, T or G) of the priming protospacer (PPS) triggered strong spacer acquisition at CRISPR locus 1 (C1), while no new spacer was detected in the negative control sample carrying the empty vector pZM15Asp. However, CC PAM, which conferred strong DNA interference (Fig. S1)^34^, triggered weak acquisition (Fig. 1c), due to a single nucleotide mutation at CC PAM (CC to CT) or protospacer sequence, or a deletion of the 3’ portion of the PPS on the plasmid. In contrast, other NN combinations could not trigger strong primed spacer acquisition (Fig. 1c). Further, CRISPR locus 2 and 3 showed similar acquisition patterns to locus 1, indicating that the leader sequences were all functional (Fig. 1c). However, acquisition efficiency at locus 2 and 3 were weaker than locus 1 (Fig. 2d), demonstrating the leader sequence variations had an effect on primed spacer acquisition (Fig. S2). Moreover, by extending the culture time, expanded bands on the gel representing new spacers became more apparent than the parental bands (Fig. 1d), and TT, GT, and TG PAM sequences triggered primed acquisition at 5 (Fig. 1d), 10, and 15 days (Fig. S3), respectively, suggesting NT or TN PAMs, in addition to CN or NC PAMs, had the ability to promote priming.

**Figure 1.**
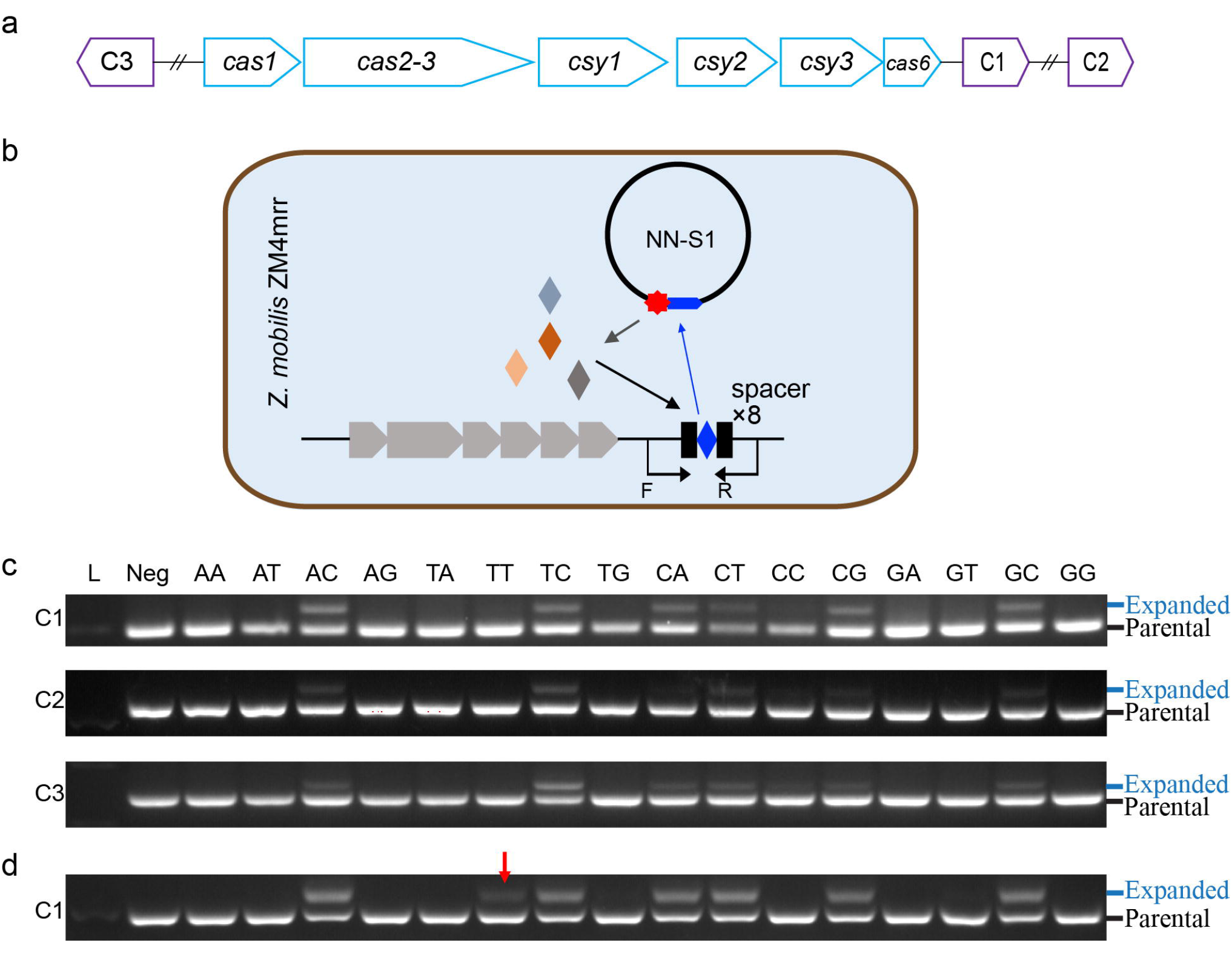
Identification of leader and PAM variants for primed spacer acquisition in *Z. mobilis* ZM4. **a**, organisation of CRISPR-Cas subtype I-F module in *Z. mobilis* ZM4. Three CRISPR arrays (C1, C2, and C3) were indicated. **b**, Schematic of the primed spacer acquisition assay. The plasmids carrying a protospacer matching spacer 1 of CRISPR locus 1 adjacent to different PAMs (NN, N = A, T, C or G) were transformed into *Z. mobilis* ZM4mrr. Three single colonies of each transformants were selected and cultured for 1 day in RM medium. The *cas* genes are indicated in grey arrows, repeats are indicated in black squares, and spacers are indicated by diamonds. **c**, Detection of spacer acquisition at three CRISPR loci (C1, C2, and C3) on 1.5% agarose gels. Each colony of transformant was cultured in liquid medium for 1 day and used for amplification of the leader proximal regions. The parental and expanded bands are indicated. These data represent three independent spacer acquisition analyses for each construct. **d**, cells used for amplification of the leader proximal region of CRISPR locus 1 were cultured in the same medium without antibiotics for 5days. For the extended culture, 1% cells were transferred into fresh medium daily. Red arrows indicate newly expanded bands identified after extended culture. Each gel is a representative of three repeated experiments using three independent single colonies for each construct.

**Figure 2.**
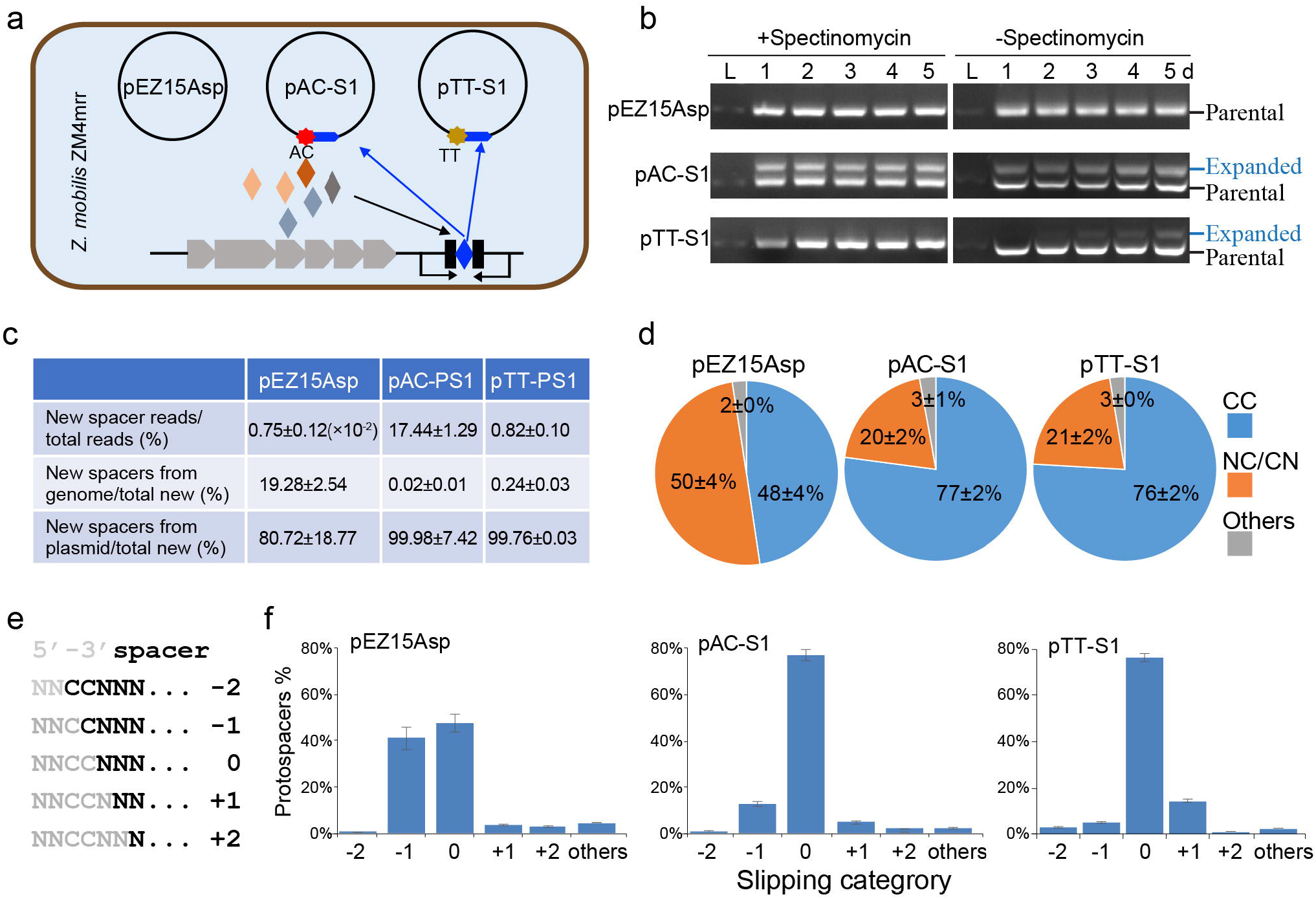
Comparison of *de novo* and primed spacer acquisition. **a**, Schematic of *de novo* and primed spacer acquisition assay. The empty plasmid (pEZ15Asp) or plasmid carrying protospacers matching a pre-existing spacer 1 of CRISPR locus 1 adjacent to two atypical PAM sequences (AC and TT) were transformed into *Z. mobili*s ZM4mrr. **b**, Detection of spacer acquisition at CRISPR locus 1 of *Z. mobilis* transformants cultured in medium with or without antibiotics on 1.5% agarose gels. Each gel is a representative of three repeated experiments using three independent single colonies for each construct. **c**, The PCR-amplified leader-proximal region containing newly adapted spacers was analysed by high-throughput sequencing, and the data are summarised. Data were collected from three independent colonies of each transformant (two independent colonies for pEZ15Asp transformant). **d**, analysis of the PAM sequences of protospacers identified in the high-throughput sequencing. **e**, schematic representation of slipping, which results in capture of spacers that map to non-canonical PAMs. Spacer sequence is indicated by black letters, and 5’-end CC PAM is indicated by (specify symbol/identifier here). **f**, frequency of slipping events observed in each dataset. Others indicated that 5’-end of protospacer was >2 nt from the CC PAM.

### Self-discrimination and PAM recognition in *de novo* and primed spacer acquisition

Cells carrying pEZ15Asp, pAC-S1, and pTT-S1 (Fig. 2a) were cultured for 5 days to analyse acquisition efficiency. Plasmid pEZ15Asp didn’t induce detectable acquisition, even with the extended cultivation time or the presence/absence of antibiotics (Fig. 2b). The pCC-S1 construct showed strong primed acquisition in ± antibiotic medium with the extended culture (Fig. 2b). The pTT-S1 plasmid only induced moderate acquisition in antibiotic-free medium (Fig. 2b). The expanded bands of day 1 antibiotic-free cultures were purified and analysed by high-throughput sequencing. The efficiencies of *de novo* spacer acquisition and pTT-S1 induced primed acquisition were ~0.04% and ~4.70% of the pAC-S1 induced primed acquisition (Fig. 2c). Most of the new spacers (>99%) were adapted from plasmid DNA in pAC-S1 and pTT-S1 primed acquisition, while only ~80% of new spacers were adapted from plasmid DNA in pEZ15Asp-induced *de novo* spacer acquisition (Fig. 2c), indicating more efficient self-discrimination during primed acquisition than *de novo* acquisition in the *Z. mobilis* subtype I-F system.

Further, all constructs recognised protospacers with a high proportion of non-canonical CC PAM. Approximately 50% of protospacers identified during *de novo* acquisition had NC/CN PAM sequences, while ~20% from both primed acquisition experiments had NC or CN PAMs (Fig. 2d), which is discordant with other studies on subtype I-F systems reporting that primed protospacers strictly required canonical PAM sequences ^33,35^. Protospacers with NC or CN PAMs could probably be recognised at the canonical CC PAM but were acquired by sliding with −2, −1, +1, or +2 nucleotides related to CC PAM (Fig. 2e and f), confirming that acquisition of these protospacer was initiated at a canonical CC PAM but not a random PAM.

### Identification of seed sequence for DNA interference and primed acquisition

Systematic transversion substitution was introduced at each nucleotide of the protospacer in the pCC-S1 construct (Fig. 3a). Single-nucleotide mutations at nucleotides 1–5 and 7 (partially) rescued transformation efficiency (Fig. 3b), indicating that these nucleotides acted as the seed sequence for CRISPR interference. Single colonies of these transformants were isolated, and sequencing results revealed no additional mutations at the PAM or protospacer sequences. However, designed mutations at this region triggered moderate primed spacer acquisition (Fig. 3c). Mutations at nucleotides 6 and 8–16 had no effect on CRISPR interference (Fig. 3b). From isolated mutant colonies, plasmids in some cells showed additional mutations at the PAM or protospacer sequences, resulting in weak primed acquisition, while the intact plasmid in construct M14 induced strong primed acquisition (Fig. 3c).

**Figure 3.**
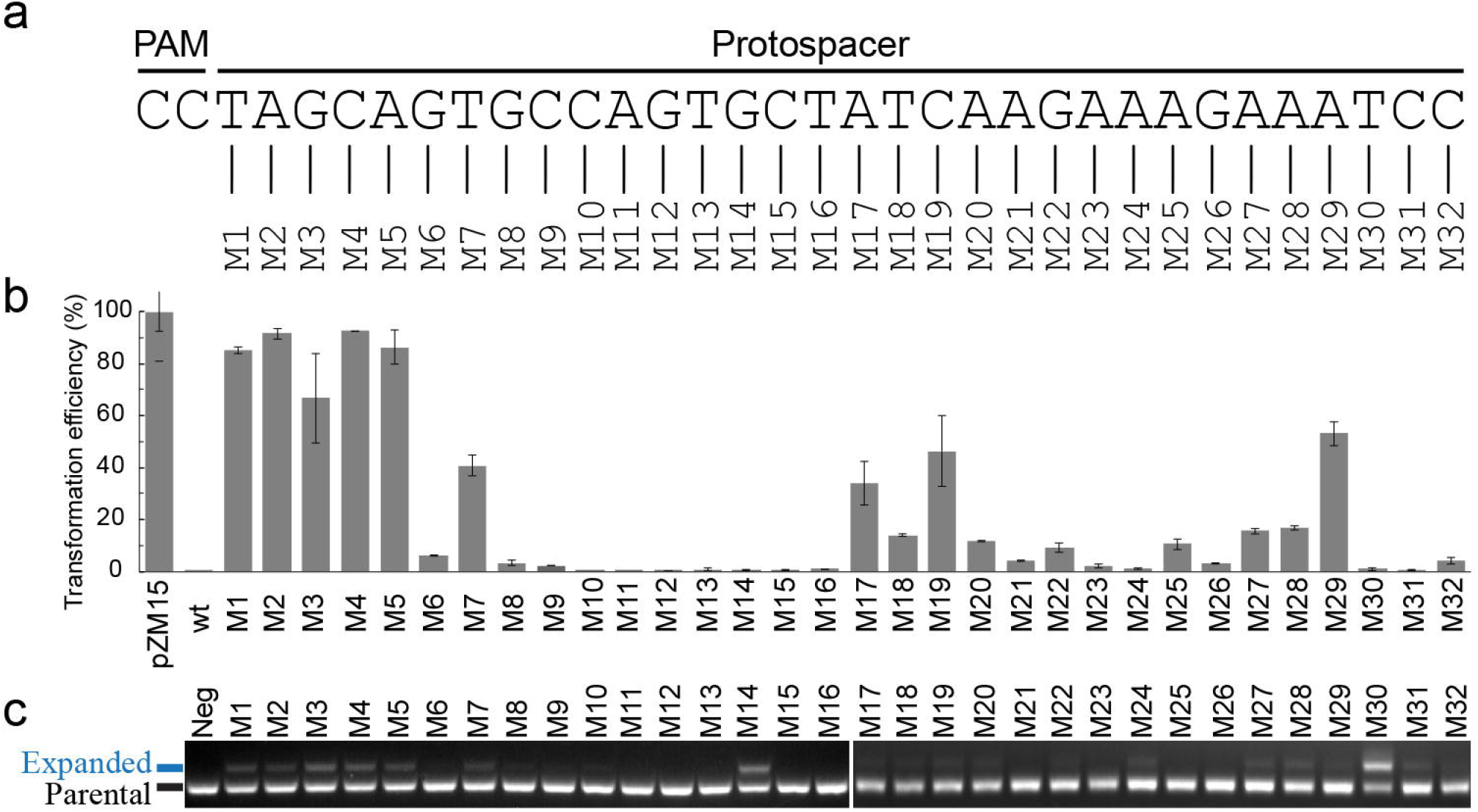
Seed sequence of the protospacer for DNA interference and primed spacer acquisition. **a**, Protospacer sequence matching spacer 1 of CRISPR locus 1 with a 5’-CC PAM was cloned into pEZ15Asp vector. A single transversion substitution was introduced at each nucleotide. **b**, each plasmid mutant was transformed into *Z. mobili*s ZM4mrr, and transformation efficiencies related to the pEZ15Asp plasmid are shown. **c**, Detection of spacer acquisition at CRISPR locus 1 on 1.5% agarose gels. Each colony of transformants was cultured in liquid medium for 1 day and used for amplification of the leader proximal regions. The parental and expanded bands are indicated. These data represent three independent spacer acquisition analyses for each construct.

Further, mutations at the region from 17–29 nucleotides were found to partially rescue transformation efficiency (Fig. 3b). Most single colonies of these transformants showed additional mutations at the protospacer sequence on the plasmid, and weak primed acquisition was found (Fig. 3c). Nucleotides 30–32 were not essential for interference (Fig. 3b). However, only two single colonies were isolated for M30, and the intact plasmid in these two colonies showed the strongest primed acquisition efficiency (Fig. 3c).

### Priming protospacer determines location and strand biases of new acquisitions

New spacers were visualised by mapping their cognate protospacer locations on the plasmids (Fig. 4). Protospacer mapping was remarkably consistent across the replicates for the colonies carrying pEZ15Asp, pAC+ and pAC-plasmids (Fig. S4). In the *de novo* acquisition experiment (colonies carrying pEZ15Asp), protospacer hotspots were identified near the origin for plasmid replication in *Z. mobilis*, however, another hotspot was identified at the negative strand on the antibiotic-resistance gene in two replicates (Fig. S4a). Primed spacer acquisitions showed distinct patterns for protospacer distributions related to the PPSs (Fig. 4a and b). A significant protospacer peak was found on the same stand of PPS around the priming sites (Fig. 4a and b). On the scale of whole plasmid DNA, primed spacer acquisition showed preference for the paired strand related to the location of the PPSs (Fig. 4c and d) and showed preference for the 3’ regions of both DNA strands related to the location the PPS (Fig. 4c and e).

**Figure 4.**
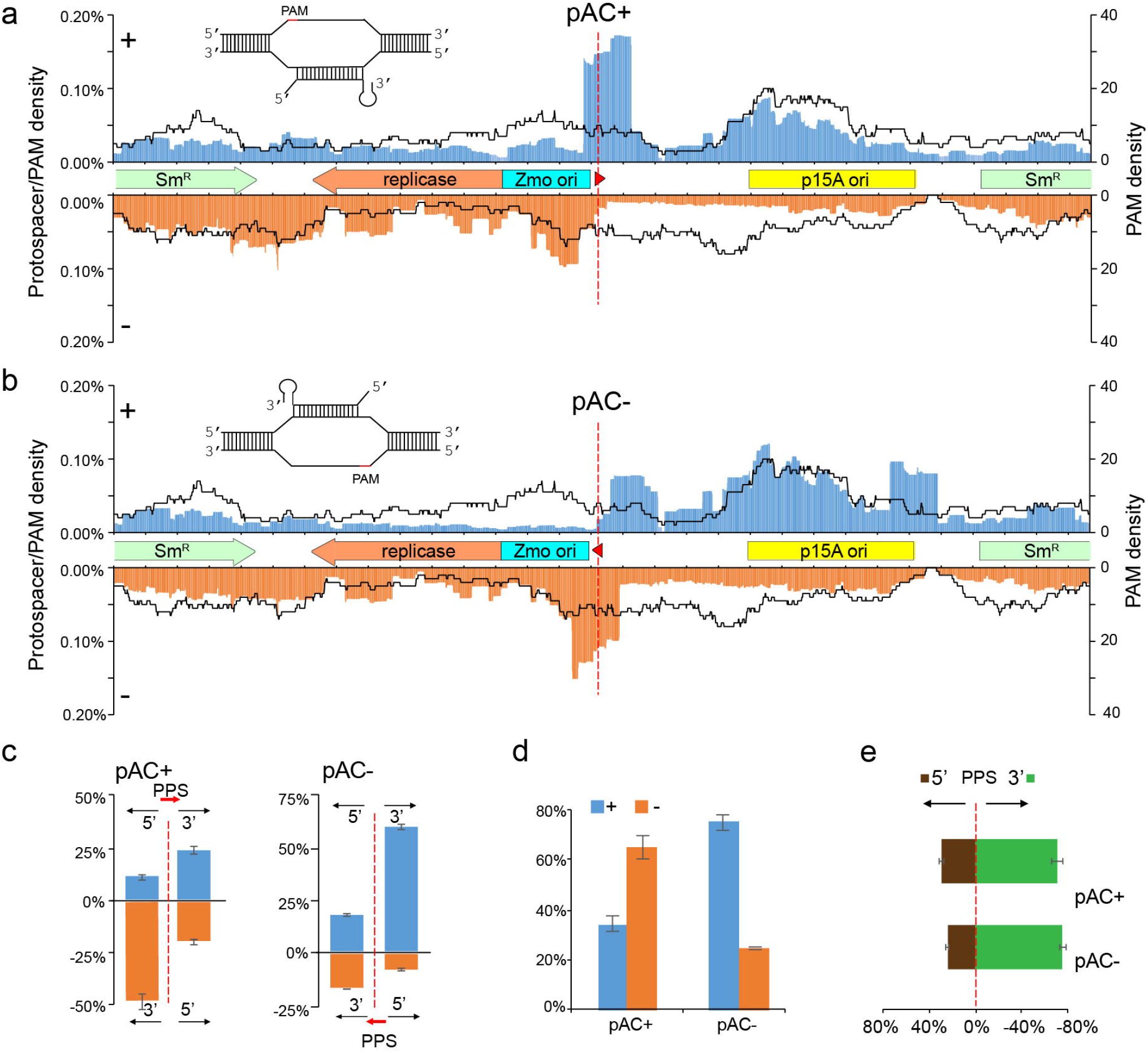
Protospacer distribution reveals biased spacer incorporation. The protospacer locations were mapped on **a**, pAC+, and **b**, pAC- plasmids using a sliding 150 nt binning window. Protospacers on the plus and minus strand are indicated by blue and orange, respectively. PAM distributions in a sliding 150 nt binning window are indicated by black lines. The position of priming protospacer (PPS) in pAC+ and pAC-plasmids is indicated with a red dash line and arrows. Plasmid-encoded genes and cis acting elements are indicated by arrows. **c**, Proportion of protospacers distribution at the 5’ and 3’ of the PPS on each strand. PPS is indicated by a red dash line, and the direction is indicated by a red arrow. Blue and orange squares indicate the plus and minus strands, respectively. **d**, proportion of protospacers present on each strand. Blue and orange squares indicate the plus and minus strands, respectively. **e**, Proportion of protospacers distribution at the 5’ and 3’ of the PPS on both strands. Data shown represent mean ± SD for the different three (for pAC+) or two (for pAC−) independent transformants of each plasmid.

### Microhomology-mediated end-joining of self-targeted DNA breaks

To study the host cell to escape targeting guided by adapted self-spacers, we constructed self-interference plasmids carrying mini-CRISPR cassette encoding spacers matching a non-essential gene (*Zmo0346*) and an putative essential folylpolyglutamate synthase gene (*Zmo0582*) (Fig. 5a). Transformation efficiency of pCC-S1 was very low, since this plasmid was targeted by the host CRISPR complex guided by the CRISPR locus 1-derived spacer (Fig. 5b). Similarly, transformation efficiency of the pS0582 plasmid, which encoded a spacer matching a protospacer on the putative essential gene *Zmo0582,* was low, indicating that the *Z. mobilis* CRISPR-Cas system cleaved its genomic DNA, leading to cell death without (or with inefficient) DNA damage repair (Fig. 5b). Surprisingly, transformation efficiency of the pS0346 plasmid, encoding a spacer matching a protospacer on the non-essential gene *Zmo0346,* was much higher than that of pS0582 (Fig. 5b). This result suggested CRISPR RNA-guided DNA double strand break probably was repaired by some mechanism with very high efficiency.

**Figure 5.**
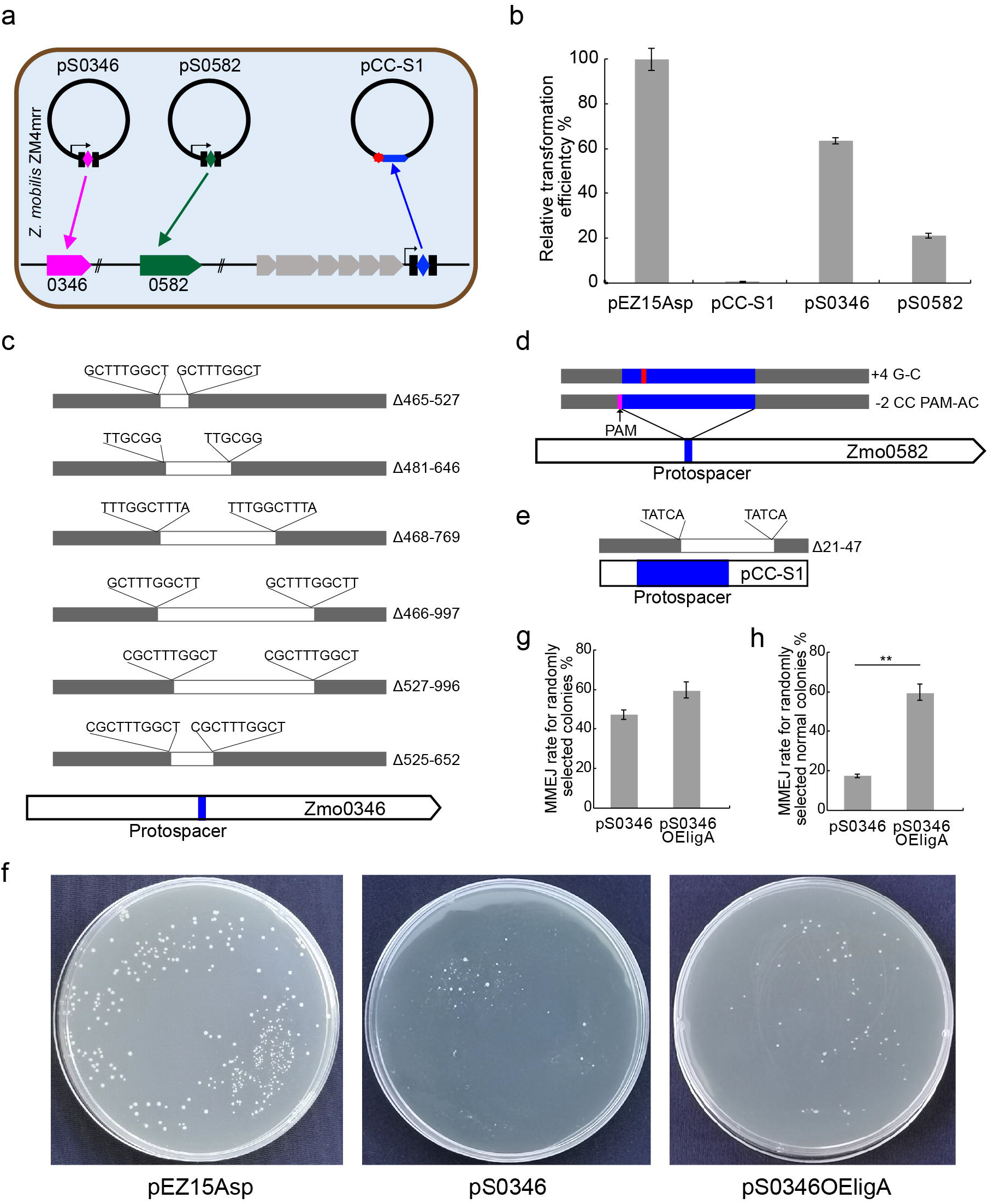
Microhomology-mediated end-joining repair of self-targeted DNA. **a**, schematic of self-targeting assay. Self-targeting plasmids carrying “Repeat-Spacer-Repeat” expression cassettes, in which the spacer matches to the Zmo0346 or Zmo0582 gene, were transformed into *Z. mobili*s ZM4mrr. The challenging plasmid pCC_PS1 in Figure 1 was used as the control. **b**, transformation efficiency of the empty plasmid, self-targeting plasmids, and challenging plasmid. **c**, **d**, and **e**, locations of mutations at the genomic Zmo0346, Zmo0582 genes, and protospacer on the plasmid in the randomly selected single colonies carrying pS0346, pS0582, or pCC-S1 plasmids. Protospacer location is indicated by a blue bar for each gene. Grey bars indicate sequenced regions, and white bars indicate deletion regions. Direct repeat sequences flanking the deletion regions are indicated upon the bars. For the Zmo0582 gene, PAM is indicated by an arrow, and a point mutation within the protospacer is indicated by a red bar. Deletion or mutation sites, related to the gene start codon for **c** or the first nucleotide of the protospacer for **d** and **e**, are indicated at the right of each graph. **f**, single colonies of *Z. mobilis* transformed with pEZ15Asp, pS0346, and pS0346OEligA plasmids on the plates. Normal and tiny colonies were indicated by arrows. MMEJ repair rates at Zmo0346 gene locus of randomly selected single colonies (**g**), or randomly selected normal colonies (**h**) carrying pS0346 or pS0346OEligA plasmids. The significance of the MMEJ efficiency was determined suing a *t*-test; p<0.01 **.

To explain the phenomenon, dozens of single colonies transformed with pS0346 and pS0582 plasmids were selected, and the target gene loci were PCR-amplified for sequencing. Different deletion mutations or point mutations were identified at the *Zmo0346* gene (Fig. 5c) and *Zmo0582* gene loci (Fig. 5d). For the essential gene *Zmo0582*, most mutations occurred at the CC PAM or the seed sequence of the protospacer (Fig. 5d), leading to host escape of self-interference. For the non-essential *Zmo0346* gene, deletion regions covering the protospacer were identified, leading to escape of CRISPR interference (Fig. 5c). Deletions at the *Zmo0346* gene locus were different in length (63-531 bp); however, short direct repeats, 6–10 bp in length, flanking the deletion regions were identified (Fig. 5c). Similar result was also found for the experiments designed to target other genes. For example, self-targeting at Zmo1062 gene resulted in ~75% randomly selected colonies was repaired with deletion mutations covering the target site (Fig. S5). We also sequenced the plasmids from the single colonies transformed with the pCC-S1 plasmid and found point mutations at the PAM sequence (20%) or deletion mutations at the protospacer sequence (~10%). Analysis of the deletion mutations on the plasmid found a 5-bp direct repeat adjacent to the deletion region (Fig. 5e), indicating that CRISPR interference at both genomic and plasmid loci induced microhomology-dependent end-joinning (MMEJ), resulting in deletion mutations.

We further studied whether the replicative NAD^+^-dependent Ligase-A (LigA) was involved in MMEJ. Overexpression *ligA* gene on the plasmid reduced the transformation efficiency (Fig. S6), suggesting cytotoxicity. Further, the *ligA* gene was overexpressed on the pS0346 self-interference plasmid (pS0346OEligA), and its end-joining efficiency was assessed. Transformants carrying the control plasmid pEZ15Asp showed normal (large) colonies on the antibiotic plates; however, transformants carrying the self-interference plasmid pS0346 showed many small colonies and few normal colonies (Fig. 5f). PCR analysis indicated that ~46% of the random selected colonies transformed with pS0346 showed MMEJ repair at the Zmo0346 locus (Fig. 5g). All transformants carrying the pS0346OEligA plasmid formed large colonies on the plates, similar to the control plasmid pEZ15Asp (Fig. 5f). PCR analysis indicated that ~60% of the randomly selected colonies overexpressing LigA showed MMEJ repair at the Zmo0346 locus. However, only ~10% of the normal colonies carrying pS0346 interference plasmid were MMEJ repaired (Fig. 5g), indicating that *ligA* overexpression increased efficiency of MMEJ repair of the self-targeted DNA break.

## Discussion

### Diversity of primed acquisition in different CRISPR subtypes

Primed spacer acquisition has been studied in different CRISPR-Cas subtypes^7,16,20,21,36^. In *Z. mobilis* subtype I-F system, more than 50% of PAM mutations promoted primed spacer acquisition (Fig. 1C and Fig. S2), similar to other systems, such as *E. coli* subtype I-E ^37^ and *H. hispanica* subtype I-B ^7^. Priming sampled protospacers mostly from the targeted plasmid in *Z. mobilis* subtype I-F system, consistent with previous studies which reported that priming reinforces immunity against invasive genetic elements ^38^. In nearly all of the studied primed acquisition systems, for example, *E. coli* subtype I-E ^16^, *H. hispanica* subtype I-B ^17^, *P. atrosepticum* subtype I-F ^33^ and *L. pneumophila* subtype I-C ^36^, primed acquisition is strictly dependent on recognition of a canonical PAM. It was found that >90% of the identified protospacers harboured a canonical PAM sequence; however, in the *Z. mobilis* subtype I-F system, only ~77% of identified protospacers had a canonical CC PAM (Fig. 2d). Further analysis of nearby sequences of protospacers with less canonical PAM, a ±2 nt regions showed identity with CC PAM, indicating a slipped PAM recognition pattern as reported previously ^33,36^, but occurring at a higher frequency. These slipped-acquired spacers might guide the CRISPR surveillance complex to recognise the NC or CN PAM (Fig. 2e), therefore triggering a second-round priming.

Priming acquired new spacers close to the PPS, and the protospacer distribution of most subtypes was strand- and direction-biased ^38^. Previous studies demonstrated that subtypes I-B, I-C, and I-E showed biases towards the acquisition of new spacers from the same strand as the PPS ^17,30,36^; however, the *P. atrosepticum* subtype I-F system acquired spacers almost equally from both strands ^21,33^. Here, priming in *Z. mobilis* subtype I-F system was biased towards acquired new spacers from the paired strand of the PPS on the whole plasmid DNA level (Fig. 4d), opposite to other systems. However, priming highly preferred the same strand around the PPS regions in Z. mobilis subtype I-F system (Fig. 4a and b). Regarding directional bias, the *P. atrosepticum* subtype I-F system showed a priming bias towards the 5’ direction ^33^. In contrast, in *Z. mobilis* subtype I-F system, priming was biased towards the 3’ direction of the PPS site (Fig. 4e), similar to the I-B, I-C, and I-E systems ^38^. Asymmetric protospacer distribution was generally found in our priming experiments (Fig. 5b–d), similar to the *P. atrosepticum* subtype I-F system ^33^. Protospacers that mapped near the PPS site showed clustering. However, a high density of protospacers at the far 3’ region of PPS was detected (Fig. 4b and c), opposite of the I-F system^21,33^, but similar to I-B, I-C, and II-A systems ^38^.

### Primed acquisition and MMEJ cooperate to avoid self-targeting

In this study, we found primed spacer acquisition was much more robust than *de novo* acquisition (Fig. 2b), showing greater specificity to invasive DNA (Fig. 2c). However, the pre-existing spacers required *de novo* acquisition for “first step” integration (Fig. 6). While, similar to other systems, *de novo* acquisition adapted a large proportion of spacers from the host genomic DNA in our system (Fig. 2c), possibly resulting in self-targeting (Fig. 6). In *Z. mobilis*, transformation of self-targeting plasmids still resulted in a relatively high transformation efficiency (Fig. 5b). Further analysis of the target locus of transformants revealed that the MMEJ pathway was involved in repair of the host DNA DSBs, rescuing the targeted host cells (Fig. 5c and Fig. 6). The MMEJ pathway is an error-prone repair process, resulting in deletion of target regions covering the protospacer site (Fig. 5c) ^39^ and finally avoiding self-targeting. Therefore, *Z. mobilis* developed primed acquisition and robust DNA repair systems to promote specific CRIPSR immunity against invasive genetic elements and avoid self-targeting (Fig. 6).

**Figure 6.**
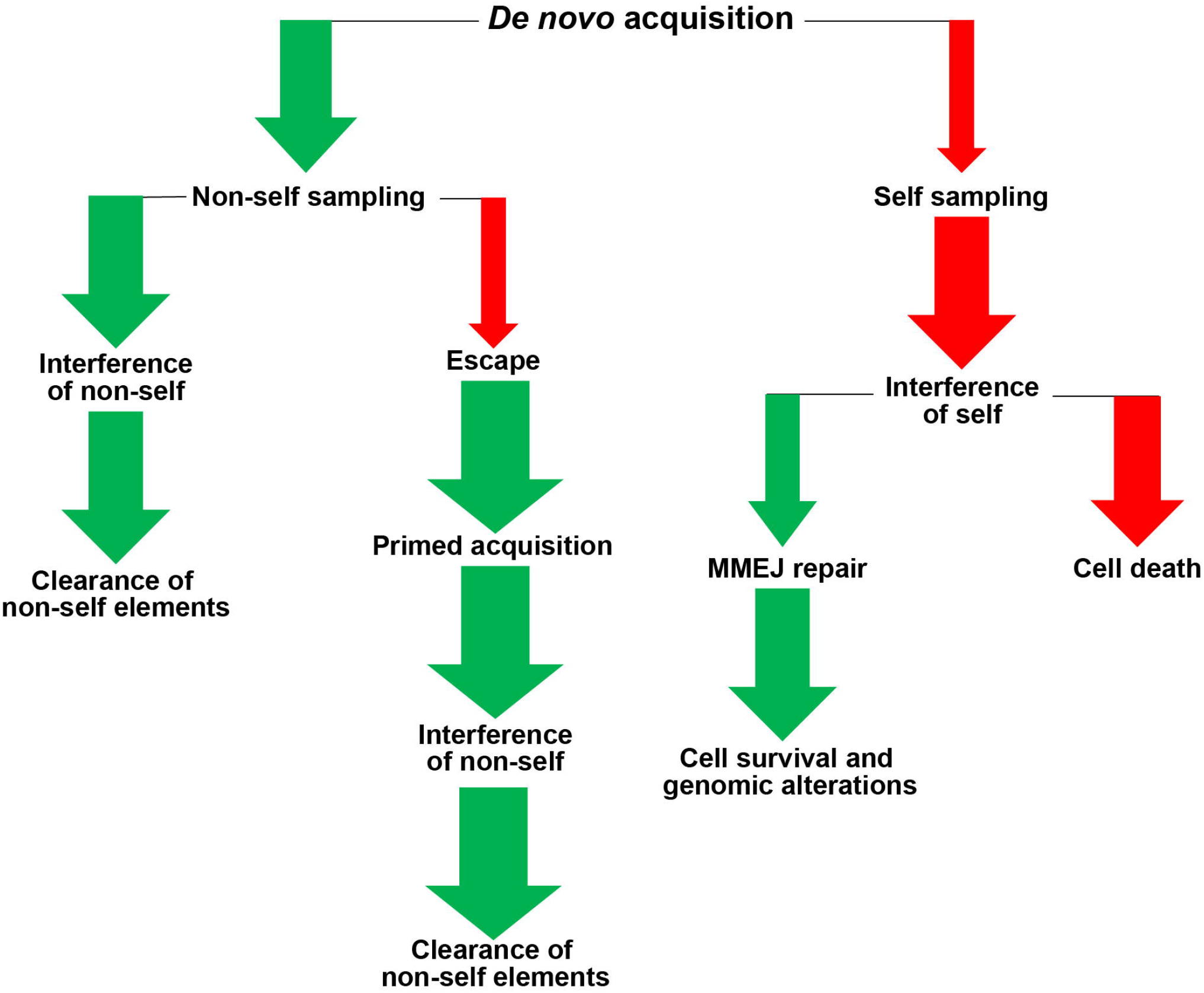
Proposed model for CRISPR immunity against invasive genetic elements in *Z. mobilis* subtype I-F system. *De novo* spacer acquisition samples spacers from both invasive genetic elements and host DNA. Non-self DNA sampling leads to CRIPSR interference of invasive genetic elements or escape of CRISPR interference. Escape triggers primed acquisition, finally leading to interference and clearance of invasive (non-self) genetic elements. Host self DNA sampling leads to interference of self DNA, resulting in cell death. However, MMEJ repairs the self-spacer-guided DNA DSBs, resulting in cell survival and genomic alterations. The width of the arrows indicates the probabilities. Red arrows indicate threats to the host, and green arrows indicate immunity against the invasive genetic elements.

MMEJ also repair the targeted plasmid DNA (Fig. 5b). Although very few single colonies were isolated, ~10% of these single colonies were found carrying a short deletion (27 bp) at the protospacer and its flanking region (Fig. 5e). In the *de novo* acquisition, protospacers on the invasive genetic elements formed a hotspot at the replication origin (Fig. 4a) ^25^, while most protospacers from the host genomic DNA were located on non-essential genes (Table S3). Therefore, MMEJ-mediated deletion of protospacers flanking the Zmo origin would probably hinder plasmid replication, while deletion of non-essential genes was nonlethal for the cells.

Error-prone repair systems contributing to avoid CRISPR self-targeting could be widespread in bacteria. For example, the non-homologous end-joining (NHEJ) pathway is co-present with the CRISPR-Cas system in some bacterial species ^40,41^. However, the NHEJ system has no effect on CRISPR immunity when both NHEJ genes and CRISPR-Cas subtype II-A system were co-expressed in a heterogeneous host ^40^. Furthermore, *Mycobacterium smegmatis* encodes subtype III-A and NHEJ systems; its NHEJ system only works for heterogenous CRISPR-FnCpf1 cleaved host DNA, not for the heterogenous CRISPR-TdCas9_m or CRISPR-NmCas9 systems ^41^. Another end-joining repair system, MMEJ, found in *E. coli* has been experimentally demonstrated to repair DNA DSBs ^39,42^. Here, the repair efficiency for DSBs by MMEJ system in *Z. mobilis* was significantly higher than that in *E. coli* (~3.0×10^−1^ vs ~3.0 ×10^−5^, Table S2) ^39,43^. In summary, our results demonstrated the function of the MMEJ system to cooperate with the CRISPR-Cas system to repair CRISPR self-targeted host DNA, therefore promoting immune specificity against invasive genetic elements.

## MATERIALS AND METHODS

### Strains, growth, and transformation of *Z. mobilis*

*Z. mobilis* strains, including ZM4, ZM4mrr^44^, and ZM4mrr carrying different plasmids, were cultured in RM medium (20 g/L glucose, 10 g/L yeast extract, and 2 g/L K_2_HPO_4_) at 37°C. *Z. mobilis* ZM4mrr lacking an endonuclease gene was used as the genetic host in this study. Plasmid DNA, modified in the *E. coli* Trans10 strain, was transformed into *Z. mobilis* ZM4mrr by electroporation, and transformants were selected on RM medium agar plates with 100 μg/mL spectamycin. Single colonies from the plates were cultured in RM liquid medium for cell growth. *E. coli* Trans10 cells used for DNA cloning were cultured at 37°C in Luria–Bertani medium, and spectamycin was added to the culture to a final concentration of 100 μg/mL, if required.

### Construction of plasmids

The PAM variable challenging (or priming) plasmids were constructed by cloning the “NN-S1” (NN = any nucleotide, S1 = the first spacer of CRISPR locus 1 of *Z. mobilis* ZM4) oligonucleotides into the shuttle vector pEZ15Asp at EcoRI/XbaI restriction sites, resulting in 16 (4^2^) challenging plasmids. The seed sequence variable priming plasmids were constructed by introduction of a transversion substitution at each nucleotide of the “Spacer 1” of the plasmid (pCC-S1) carrying the “CC-Spacer 1” cassette. The interference plasmids carrying the expression cassette of “Repeat-Spacer-Repeat” against genomic loci, Zmo0346 and Zmo0582 genes, were constructed by cloning the 32-bp sequences following a CC PAM on these gene coding sequences into the pEZ15Asp vector under control of the *cas1* gene promoter (P*cas1*), resulting in interference plasmid pS0346 and pS0582. Zmo0364 (*ligA* gene) overexpression plasmids were constructed by cloning the Zmo0364 open reading frame under control of the Zmo0367 promoter ^45^ into pEZ15Asp or the interference plasmid pS0346.

### PCR amplification of the leader-proximal CRISPR regions

*Z. mobilis* strains harbouring an empty vector (pEZ15Asp) or the challenging (priming) plasmids were cultured in 10 mL of RM medium at 37°C without shaking for one day or passaged daily for 15 days by transferring 100 μL cell culture to 10 mL fresh medium with or without antibiotics. Samples of each culture (0.1 mL) were taken at a desired time point, and total DNA from these cells was used as the PCR template. The leader-proximal regions of three CRISPR loci or only CRISPR locus 1 were amplified by PCR using Taq polymerase and the forward/reverse primer sets (Table S1) for locus 1, 2, or 3, respectively. PCR products were separated on a 1.5% agarose gel and visualised by ethidium bromide staining.

### High-throughput sequencing and bioinformatics analysis

Three single colonies of each transformant were selected from the plate and inoculated in RM medium for growth. Genomic DNA was extracted from each enlarged culture of these single colonies and used for amplification of the leader-proximal region of CRISPR locus 1. Sequencing was carried out as described previously ^46^. Using the BLASTN program against pEZ15Asp plasmid or pEZ15Asp carrying PPS or the *Z. mobilis* ZM4 genome sequence ^47^, the protospacer sequences were identified for new integrated spacers. The 2-bp sequence at the 5’-end of each protospacer was considered as the PAM region. Perl scripts were run to analyse the protospacers. All high-throughput sequencing reads have been deposited into the SRA database with the accession number PRJNA579098. The protospacer locations of the newly acquired spacers in the CRISRP locus from *de novo* acquisition or primed acquisition were mapped on pEZ15Asp and the priming plasmids pAC-SS1(+) and pAC-S1(−) (pAC+ and pAC- for short), respectively, using a sliding 150 nt binning window, as described previously ^33^.

## Acknowledgements

This work was supported by the National Natural Science Foundation of China (No. 31671291 to N.P., No. 31570055 to M.H. and 31801035 to Y.L.), and the Fundamental Research Funds for the Central Universities (No. 2662019PY028 to N.P.). Funding for open access charge: National Natural Science Foundation of China (No. 31671291).

## Author contributions

X. W. and B. W. conducted the experiments, Z. Z., T. L. and Y. L. analyzed the acquisition data, M. H., G.H. and N. P. designed the experiments and analyzed the data. N.P. wrote the manuscript.

