## Supplementary Figure 1 for "Primed acquisition and microhomology-mediated end-joining cooperate to confer specific CRISPR immunity against invasive genetic elements"

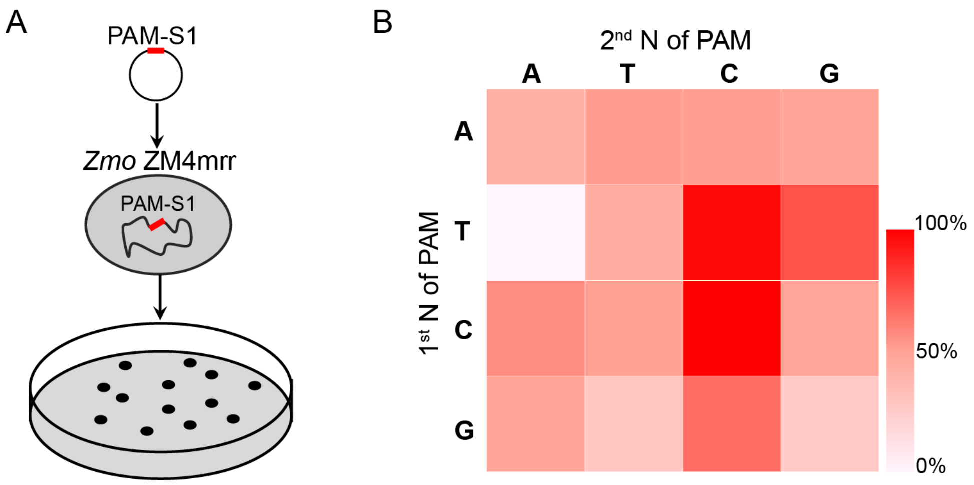


**Supplementary Figure 1. Identification of PAM variants for DNA interference.** **a,** Schematic of the plasmid interference assay. The empty shuttle vector (pEZ15Asp) or the pNN-S1 (N = A, T, C or G) vector, carrying the protospacer matching spacer 1 of CRISPR locus 1 with 16 (NN) different PAMs, was transformed into *Z. mobilis* ZM4mrr cells. **b**, Heat map of the interference efficiency. The interference efficiency was calculated as "transformation efficiency of plasmids with different PAM/transformation efficiency of the empty shuttle vector". The colour bar corresponding to interference efficiency is shown on the right. These data represent three independent plasmid interference analyses.
