## Supplementary Figure 2 for "Primed acquisition and microhomology-mediated end-joining cooperate to confer specific CRISPR immunity against invasive genetic elements"

```
A
L1 ATTTGACCCTTATTTTGACCCTCTTTTTTCGAGGGTATAAAAAATCCTTTCGATTCAATA  -91
L2 TTTTGACCCTTTATTTGACCCTCTTTTTTTGGCATGTAAAAAAATCCTTTAAAATCAATA  -91
L3 TTTTTGCCCCTAATTTGACCCTCTTTTTTCGAGACATTTAAAAATCTTTTAAAATCAACA  -91
    ***  *** *  **************** *        ******* ***  * **** *

L1 TGTTACATATGAGCATATTTTTTAGGGTTATTTTGCCTTTTTGGCGAGATATCCCTTTAT  -30
L2 GGTTAAAAATAGGCTCTTTTGGCATGTAAAAAAATCCTTTAAAATCAATAGGTTAAAAAT  -30
L3 ACTTAAAAGCAAGCCTTTTCGAGACATTTAAAAATCTTTTAAAATCAACAACTTAAAAGC  -30
     *** *     **   **     *     *     * ***      *

L1 TTTAGGGGCAATTCTATCTTTTGCCTCTA -1
L2 AGGCTCGGGATTTTTAATTATTTACTCTA -1
L3 AAGCCTGAGATTTCTATAAATCTCTTCTA -1
         *  * ** **    *    ****

B
R1 GTTCACTGCCGCACAGGCAGCTTAGAAA
R2 GTTCACTGCCGCACAGGCAGCTTAGAAA
R3 GTTCACTGCCGCATAGGCAGCTTAGAAA
   ************* **************
```

**Supplementary Figure 2. Leader and repeat sequences of *Z. mobilis* ZM4 subtype I-F CRISPR-Cas system. a**, Alignment of leader sequences and **b**, repeat sequences of three CRISPR loci of *Z. mobilis* ZM4. Highly conserved regions were boxed.
