## Supplementary Figure 3 for "Primed acquisition and microhomology-mediated end-joining cooperate to confer specific CRISPR immunity against invasive genetic elements"

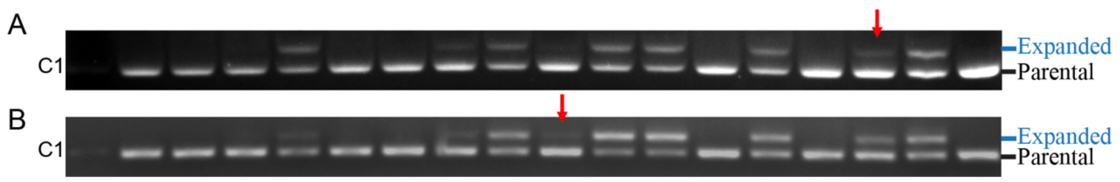


**Supplementary Figure 3. Detecting priming at CRISPR locus 1 with extended culture time for 10 (A) and 15 (B) days.** For the extended culture, 1% cells were transferred into fresh medium daily. Red arrows indicate newly expanded bands identified after extended culture. Each gel is a representative of three repeated experiments using three independent single colonies for each construct.
