## Supplementary Figure 4 for "Primed acquisition and microhomology-mediated end-joining cooperate to confer specific CRISPR immunity against invasive genetic elements"

A

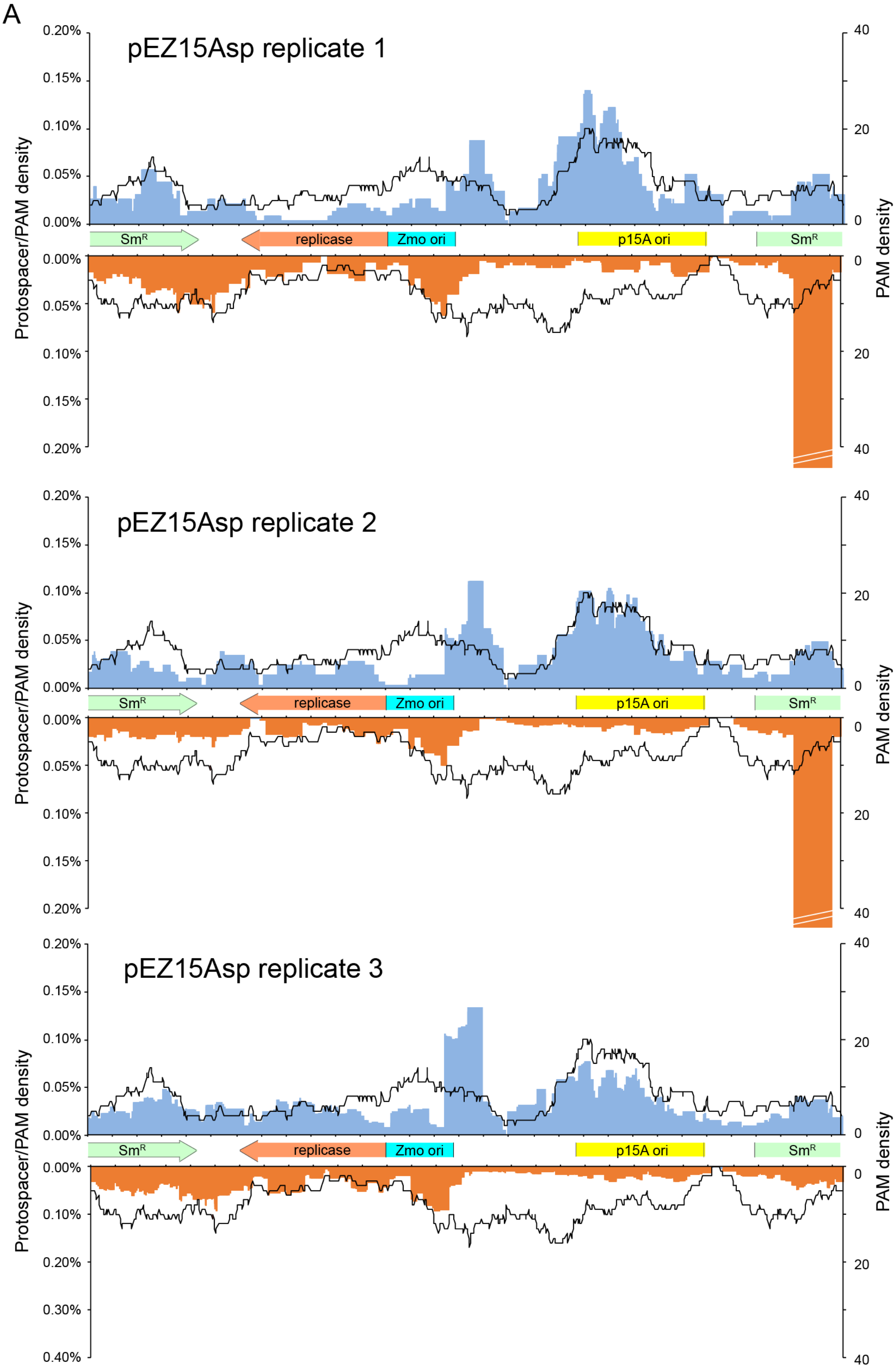
pEZ15Asp replicate 1
pEZ15Asp replicate 2
pEZ15Asp replicate 3
Protospacer/PAM density
PAM density
0.00%
0.05%
0.10%
0.15%
0.20%
0.30%
0.40%
0
20
40
SmR
replicase
Zmo ori
p15A ori

B

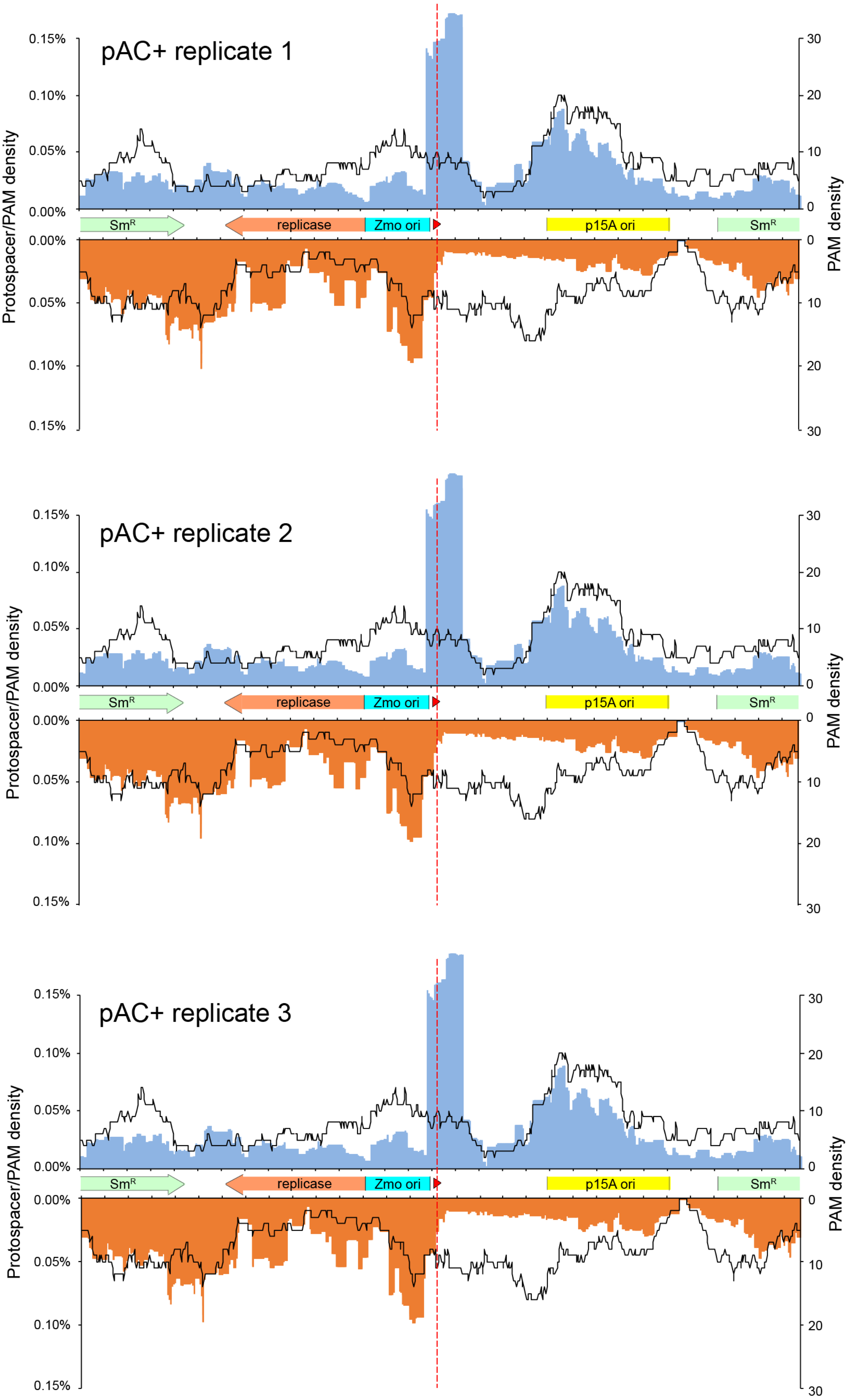
pAC+ replicate 1
0.15%
0.10%
0.05%
0.00%
0.00%
0.05%
0.10%
0.15%
Protospacer/PAM density
30
20
10
0
0
10
20
30
PAM density
SmR
replicase
Zmo ori
p15A ori
SmR
pAC+ replicate 2
0.15%
0.10%
0.05%
0.00%
0.00%
0.05%
0.10%
0.15%
Protospacer/PAM density
30
20
10
0
0
10
20
30
PAM density
SmR
replicase
Zmo ori
p15A ori
SmR
pAC+ replicate 3
0.15%
0.10%
0.05%
0.00%
0.00%
0.05%
0.10%
0.15%
Protospacer/PAM density
30
20
10
0
0
10
20
30
PAM density
SmR
replicase
Zmo ori
p15A ori
SmR

C

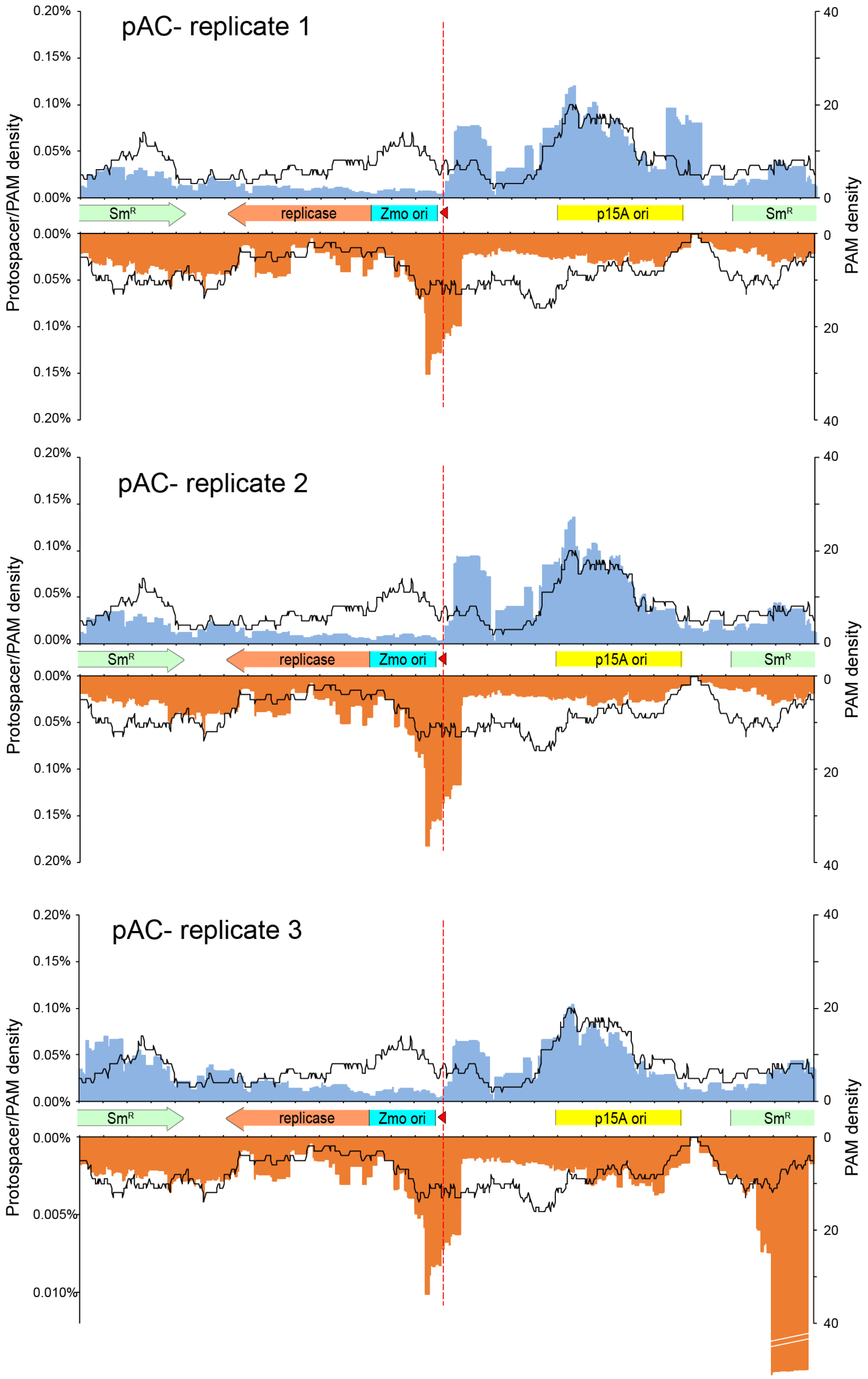


**Supplementary Figure 4. Protospacer distribution of three independent colonies carrying pEZ15Asp, pAC+ and pAC- plsamids.** The protospacer locations were mapped on **A**, pEZ15Asp, **B**, pAC+ replicates and **C**, pAC- plasmids using a sliding

150 nt binning window. Protospacers on the plus and minus strand are indicated in blue and orange respectively. PAM distributions in a sliding 150 nt binning window are indicated by the black lines. The position of PPS in pAC+ and pAC- plasmids is indicated with a red dash line and arrows. Plasmid encoded genes and cis acting elements were indicated by arrows. A strong hotspot was identified for pEZ15Asp replicate 1, 2 and pAC- replicate 3 at the same site on $Sm^R$ gene locus.
