## Supplementary Figure 6 for "Primed acquisition and microhomology-mediated end-joining cooperate to confer specific CRISPR immunity against invasive genetic elements"

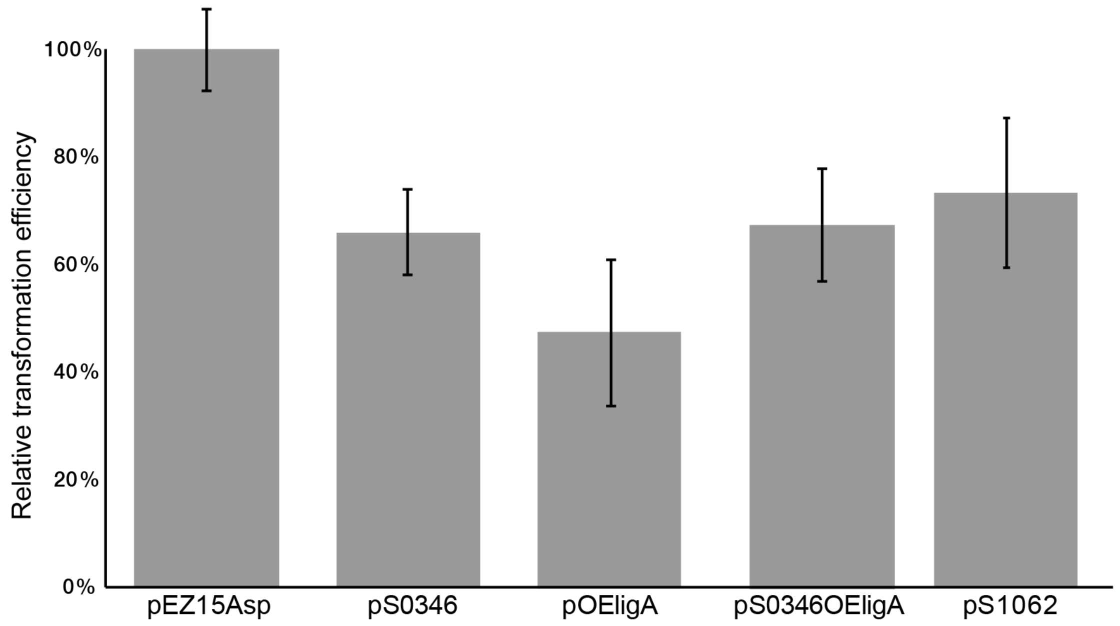


**Supplementary Figure 6. Relative transformation efficiencies of different constructs.** The plasmids pS0346 and pS1062 carried mini-CRISPR to target Zmo0346 and Zmo1062 genes. Plasmid pOEligA carried *ligA* gene overexpression cassette. Plasmid pS0346OEligA carried both self-target and *ligA* overexpression cassette.
