## Supplementary Table 1 for "Primed acquisition and microhomology-mediated end-joining cooperate to confer specific CRISPR immunity against invasive genetic elements"

Supplementary Table 1: primers used in this study.

| **Primers and nucleotides** | **Sequence (5′ to 3′)** |
|---|---|
| **Primer for cloning** | |
| PAM-S1-F-EcoRI | AATTCNNTAGCAGTGCCAGTGCTATCAAGAAAGAAATCCT |
| PAM-S1-R-XbaI | CTAGAGGATTTCTTTCTTGATAGCACTGGCACTGCTANNG |
| AC(-)S1-F-EcoRI | AATTCGGATTTCTTTCTTGATAGCACTGGCACTGCTAGTT |
| AC(-)S1-F-XbaI | CTAGAACTAGCAGTGCCAGTGCTATCAAGAAAGAAATCCG |
| SeedM1-F- EcoRI | AATTCCCGAGCAGTGCCAGTGCTATCAAGAAAGAAATCCT |
| SeedM1-R-XbaI | CTAGAGGATTTCTTTCTTGATAGCACTGGCACTGCTCGGG |
| SeedM2-F- EcoRI | AATTCCCTCGCAGTGCCAGTGCTATCAAGAAAGAAATCCT |
| SeedM2-R-XbaI | CTAGAGGATTTCTTTCTTGATAGCACTGGCACTGCGAGGG |
| SeedM3-F- EcoRI | AATTCCCTATCAGTGCCAGTGCTATCAAGAAAGAAATCCT |
| SeedM3-R-XbaI | CTAGAGGATTTCTTTCTTGATAGCACTGGCACTGATAGGG |
| SeedM4-F- EcoRI | AATTCCCTAGAAGTGCCAGTGCTATCAAGAAAGAAATCCT |
| SeedM4-R-XbaI | CTAGAGGATTTCTTTCTTGATAGCACTGGCACTTCTAGGG |
| SeedM5-F- EcoRI | AATTCCCTAGCCGTGCCAGTGCTATCAAGAAAGAAATCCT |
| SeedM5-R-XbaI | CTAGAGGATTTCTTTCTTGATAGCACTGGCACGGCTAGGG |
| SeedM6-F- EcoRI | AATTCCCTAGCATTGCCAGTGCTATCAAGAAAGAAATCCT |
| SeedM6-R-XbaI | CTAGAGGATTTCTTTCTTGATAGCACTGGCAATGCTAGGG |
| SeedM7-F- EcoRI | AATTCCCTAGCAGGGCCAGTGCTATCAAGAAAGAAATCCT |
| SeedM7-R-XbaI | CTAGAGGATTTCTTTCTTGATAGCACTGGCCCTGCTAGGG |
| SeedM8-F- EcoRI | AATTCCCTAGCAGTTCCAGTGCTATCAAGAAAGAAATCCT |
| SeedM8-R-XbaI | CTAGAGGATTTCTTTCTTGATAGCACTGGAACTGCTAGGG |
| SeedM9-F- EcoRI | AATTCCCTAGCAGTGACAGTGCTATCAAGAAAGAAATCCT |
| SeedM9-R-XbaI | CTAGAGGATTTCTTTCTTGATAGCACTGTCACTGCTAGGG |
| SeedM10-F- EcoRI | AATTCCCTAGCAGTGCAAGTGCTATCAAGAAAGAAATCCT |
| SeedM10-R-XbaI | CTAGAGGATTTCTTTCTTGATAGCACTTGCACTGCTAGGG |
| SeedM11-F- EcoRI | AATTCCCTAGCAGTGCCCGTGCTATCAAGAAAGAAATCCT |
| SeedM11-R-XbaI | CTAGAGGATTTCTTTCTTGATAGCACGGGCACTGCTAGGG |
| SeedM12-F- EcoRI | AATTCCCTAGCAGTGCCATTGCTATCAAGAAAGAAATCCT |
| SeedM12-R-XbaI | CTAGAGGATTTCTTTCTTGATAGCAATGGCACTGCTAGGG |
| SeedM13-F- EcoRI | AATTCCCTAGCAGTGCCAGGGCTATCAAGAAAGAAATCCT |
| SeedM13-R-XbaI | CTAGAGGATTTCTTTCTTGATAGCCCTGGCACTGCTAGGG |
| SeedM14-F- EcoRI | AATTCCCTAGCAGTGCCAGTTCTATCAAGAAAGAAATCCT |
| SeedM14-R-XbaI | CTAGAGGATTTCTTTCTTGATAGAACTGGCACTGCTAGGG |
| SeedM15-F- EcoRI | AATTCCCTAGCAGTGCCAGTGATATCAAGAAAGAAATCCT |
| SeedM15-R-XbaI | CTAGAGGATTTCTTTCTTGATATCACTGGCACTGCTAGGG |
| SeedM16-F- EcoRI | AATTCCCTAGCAGTGCCAGTGCGATCAAGAAAGAAATCCT |
| SeedM16-R-XbaI | CTAGAGGATTTCTTTCTTGATCGCACTGGCACTGCTAGGG |
| SeedM17-F- EcoRI | AATTCCCTAGCAGTGCCAGTGCTCTCAAGAAAGAAATCCT |
| SeedM17-R-XbaI | CTAGAGGATTTCTTTCTTGAGAGCACTGGCACTGCTAGGG |
| SeedM18-F- EcoRI | AATTCCCTAGCAGTGCCAGTGCTAGCAAGAAAGAAATCCT |

| | |
|---|---|
| SeedM18-R-XbaI | CTAGAGGATTTCTTTCTTGCTAGCACTGGCACTGCTAGGG |
| SeedM19-F- EcoRI | AATTCCCTAGCAGTGCCAGTGCTATAAAGAAAGAAATCCT |
| SeedM19-R-XbaI | CTAGAGGATTTCTTTCTTTATAGCACTGGCACTGCTAGGG |
| SeedM20-F- EcoRI | AATTCCCTAGCAGTGCCAGTGCTATCCAGAAAGAAATCCT |
| SeedM20-R-XbaI | CTAGAGGATTTCTTTCTGGATAGCACTGGCACTGCTAGGG |
| SeedM21-F- EcoRI | AATTCCCTAGCAGTGCCAGTGCTATCACGAAAGAAATCCT |
| SeedM21-R-XbaI | CTAGAGGATTTCTTTCGTGATAGCACTGGCACTGCTAGGG |
| SeedM22-F- EcoRI | AATTCCCTAGCAGTGCCAGTGCTATCAATAAAGAAATCCT |
| SeedM22-R-XbaI | CTAGAGGATTTCTTTATTGATAGCACTGGCACTGCTAGGG |
| SeedM23-F- EcoRI | AATTCCCTAGCAGTGCCAGTGCTATCAAGCAAGAAATCCT |
| SeedM23-R-XbaI | CTAGAGGATTTCTTGCTTGATAGCACTGGCACTGCTAGGG |
| SeedM24-F- EcoRI | AATTCCCTAGCAGTGCCAGTGCTATCAAGACAGAAATCCT |
| SeedM24-R-XbaI | CTAGAGGATTTCTGTCTTGATAGCACTGGCACTGCTAGGG |
| SeedM25-F- EcoRI | AATTCCCTAGCAGTGCCAGTGCTATCAAGAACGAAATCCT |
| SeedM25-R-XbaI | CTAGAGGATTTCGTTCTTGATAGCACTGGCACTGCTAGGG |
| SeedM26-F- EcoRI | AATTCCCTAGCAGTGCCAGTGCTATCAAGAAATAAATCCT |
| SeedM26-R-XbaI | CTAGAGGATTTATTTCTTGATAGCACTGGCACTGCTAGGG |
| SeedM27-F- EcoRI | AATTCCCTAGCAGTGCCAGTGCTATCAAGAAAGCAATCCT |
| SeedM27-R-XbaI | CTAGAGGATTGCTTTCTTGATAGCACTGGCACTGCTAGGG |
| SeedM28-F- EcoRI | AATTCCCTAGCAGTGCCAGTGCTATCAAGAAAGACATCCT |
| SeedM28-R-XbaI | CTAGAGGATGTCTTTCTTGATAGCACTGGCACTGCTAGGG |
| SeedM29-F- EcoRI | AATTCCCTAGCAGTGCCAGTGCTATCAAGAAAGAACTCCT |
| SeedM29-R-XbaI | CTAGAGGAGTTCTTTCTTGATAGCACTGGCACTGCTAGGG |
| SeedM30-F- EcoRI | AATTCCCTAGCAGTGCCAGTGCTATCAAGAAAGAAAGCCT |
| SeedM30-R-XbaI | CTAGAGGCTTTCTTTCTTGATAGCACTGGCACTGCTAGGG |
| SeedM31-F- EcoRI | AATTCCCTAGCAGTGCCAGTGCTATCAAGAAAGAAATACT |
| SeedM31-R-XbaI | CTAGAGTATTTCTTTCTTGATAGCACTGGCACTGCTAGGG |
| SeedM32-F- EcoRI | AATTCCCTAGCAGTGCCAGTGCTATCAAGAAAGAAATCAT |
| SeedM32-R-XbaI | CTAGATGATTTCTTTCTTGATAGCACTGGCACTGCTAGGG |
| **Primers for mini-CRISPR expression cassette** | |
| Pcas1-F-EcoRI | CGGAATTCATTTACGATTGCTCGTCCTAAATAAATAAG |
| Pcas1-R | CGGGGTACCAGAAAATCTTCTGTATCTACAATGG |
| Mini-CRISPR-F | CGGGGTACCGTTCACTGCCGCACAGGCAGCTTAGAAAAGAGACCGACGTCGGTCTCAGT |
| Mini-CRISPR-R | CGGGATCCTTTCTAAGCTGCCTGTGCGGCAGTGAACTGAGACCGACGTCGGTCTCTTTT |
| TerZmo0724-F | CGGGATCCTCGAACGCGCCGAATAAGTA |
| TerZmo0724-R-SpeI | CGACTAGTATAGCCGGTAATTTTTTTGCC |
| **For host DNA interference and deletion** | |
| 0346Spacer-F | GAAAGCTTTGGCTAGTCTTGGTTTATTTGCCGCTCG |
| 0346Spacer-R | GAACCGAGCGGCAAATAAACCAAGACTAGCCAAAGC |
| 0346Larm-F-SalI | CGCGTCGACATGCAGAGATTTTCATTAAAGG |
| 0346Larm-R | CTCGGTTGCGATCCAGACAAGCCCGAAATTTAATTCTTTC |

| | |
|---|---|
| 0346Rarm-F | GAAAGAATTAAATTTCGGGCTTGTCTGGATCGCAACCGAG |
| 0346Rarm-R-SpeI | GACTAGTTTATAGCGAGTTCATATTATAAGCCC |
| 0582Spacer-F | GAAACAAGCTGGGCAATCCCGCATACAATCGGATTG |
| 0582Spacer-R | GAACCAATCCGATTGTATGCGGGATTGCCCAGCTTG |
| 1062Spacer-F | GAAAAATGACGGTTTACAGATAAGCGTTCTTGGCCT |
| 1062Spacer-R | GAACAGGCCAAGAACGCTTATCTGTAAACCGTCATT |
| **For ligA (Zmo0364) overexpression** | |
| P0367F-EcoRI | CGGAATTCCTCCAGCATTGAATCAGACTT |
| P0367R-XbaI | GCTCTAGATATTCTCGTCCTTAAAACAGAGGCC |
| ligA-F-XbaI | GCTCTAGAATGATGCGGATATTGACC |
| ligA-R-KpnI | CGGGGTACCTTAGATTTTTATACTGTCTTGCCCG |
| TerCsy4-F-BamHI | CGGGATCCTCCTTTTAAAAAATTTGTAAAATATGCTTG |
| TerCsy4-R-SalI | CGCGTCGACTGGCCAGTATTTTTACTGACATTCAT |
| **Primers for amplification of the leader proximal CRISPR regions** | |
| Leader1-F | GGGTTATTTGCCTTTTTGGCGA |
| CRISPR1S3-R | GCGCACCTTCCGGTGTCTAT |
| Leader2-F | AGATCACGATCTGTTTTTGACATC |
| CRISPR2-R | TGAAAGCATATCGTCACGCTCACAGCGT |
| leader3-F | GGCATAATATCCCTTTATTTAGGGAGA |
| CRISPR3-R | GCCATAAAATGCTCCTGCCG |

Restriction sites and mutation sites are underlined.
