## Supplementary Table 2 for "Primed acquisition and microhomology-mediated end-joining cooperate to confer specific CRISPR immunity against invasive genetic elements"

**Supplementary Table 2. Comparison of MMEJ efficiency in *E. coli* and *Z. mobilis*.**

| | *E. coli* [1,2] | *Z. mobilis* |
|---|---|---|
| [1]Positive rate (%) | ~95-98 | ~46-75 |
| [2]Editing efficiency | $\sim3\times10^{-5}$ | $\sim3\times10^{-1}$ |

[1]Positive rate: calculated by the proportion of MMEJ repaired colonies to the total randomly selected number of colonies;

[2]Editing efficiency: calculated by the proportion of MMEJ repaired colonies in the experimental group to the total number of colonies in the control group.

1 Huang, C. *et al.* CRISPR-Cas9-assisted native end-joining editing offers a simple strategy for efficient genetic engineering in Escherichia coli. *Appl Microbiol Biotechnol* **103**, 8497-8509, doi:10.1007/s00253-019-10104-w (2019).

2 Chayot, R., Montagne, B., Mazel, D. & Ricchetti, M. An end-joining repair mechanism in Escherichia coli. *Proc Natl Acad Sci U S A* **107**, 2141-2146, doi:10.1073/pnas.0906355107 (2010).
