## Supplementary Figure 5 for "Primed acquisition and microhomology-mediated end-joining cooperate to confer specific CRISPR immunity against invasive genetic elements"

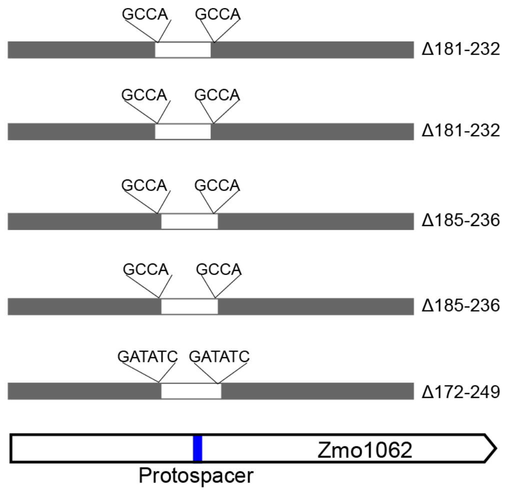


**Supplementary Figure 5. MMEJ repair at Zmo1062 gene when *Z. mobilis* ZM4mrr was transformed with a plasmid carrying the mini-CRISPR to target Zmo1062 gene.** Locations of deletion mutations at the genomic Zmo1062 in the 8 randomly selected single colonies carrying the plasmid targeting Zmo1062 gene. Protospacer location is indicated by a blue bar. Grey bars indicate sequenced regions, and white bars indicate deletion regions. Direct repeat sequences flanking the deletion regions are indicated upon the bars. Deletion sites, related to the gene start codon, are indicated at the right of each graph.
